# TomAP-MS: an improved tomato lectin affinity purification-based mass spectrometry workflow enabling ultra-deep plasma proteomics

**DOI:** 10.64898/2026.04.28.721243

**Authors:** Yusei Okuda, Hiromasa Mitsui, Ryo Konno, Daisuke Nakajima, Naho Ueyama, Osamu Ohara, Yusuke Kawashima

## Abstract

Plasma proteomics is increasingly important for biomarker discovery and disease stratification; however, comprehensive and high-throughput analysis remains challenging because of the extreme dynamic range of plasma proteins. We previously established tomato lectin affinity purification-based mass spectrometry (TomAP-MS), a workflow that enhances plasma proteome coverage via tomato lectin-mediated enrichment. The initial workflow depended on a 4% sodium dodecyl sulfate (SDS) elution, followed by SP3-based purification and digestion, which raised complexity and restricted throughput. In this study, we developed an improved TomAP-MS workflow incorporating lauryl maltose neopentyl glycol (LMNG)-assisted acid elution (LAcE), in which proteins are eluted under acidic conditions in the presence of LMNG. This process is followed by pH adjustment and direct tryptic digestion without SP3 cleanup. Compared with conventional acid elution and the original SDS/SP3 workflow, LAcE increased protein identifications while simplifying sample preparation and improving throughput. Using the optimized workflow, we identified more than 7,500 proteins from human plasma and demonstrated broader applicability in extracellular vesicle enrichment and protein interaction analysis workflows. We demonstrated that ethylenediaminetetraacetic acid plasma was the preferred specimen type, enabling the identification of over 5,000 proteins from just 1 µL of plasma, with minimal impact on proteomic profiles after up to three freeze-thaw cycles. Additionally, the analysis of plasma from 200 healthy individuals reproducibly detected 4,117 proteins across all samples, including many proteins associated with inherited disorders. These findings establish TomAP-MS with LAcE as a practical platform for deep plasma proteomics, supporting its future application in proteomics-based screening and diagnostics.

## Introduction

Plasma proteomics has become increasingly important for biomarker discovery, disease stratification, and the investigation of systemic physiological and pathological states (1, 2). Plasma obtained minimally invasively reflects various biological processes in the body, making it an appealing specimen for basic and translational research. Plasma is ideal for clinical studies, large-scale cohort analyses, and longitudinal monitoring, where repeated sampling and broad molecular readouts are essential (2, 3). Plasma’s characteristics have rendered it one of the most extensively researched biofluids in the quest for clinically significant molecular markers (1, 4). Plasma proteome profiling is commonly conducted using affinity-based platforms such as SomaScan and Olink, which enable high-sensitivity, high-throughput measurement of large panels of proteins (5, 6). These technologies have substantially accelerated biomarker discovery and population-scale protein profiling by providing robust and scalable assays (2, 5, 6). However, because they rely on predefined probe or antibody sets, they are inherently constrained to known targets and are less suited to the detection of unanticipated molecular changes (2, 7). This targeted nature also limits their ability to comprehensively characterize the plasma proteome in a truly hypothesis-free manner (1).

In contrast, liquid chromatography–mass spectrometry (LC–MS)-based plasma proteomics enables untargeted and direct protein measurement (8). This approach has continued to advance through improvements in sample preparation, peptide separation, mass spectrometric acquisition, and computational analysis (3). These developments markedly enhanced the analytical depth, reproducibility, and utility of LC–MS-based plasma proteomics (9, 10). Moreover, novel enrichment technologies, including nanoparticle-based platforms such as those developed by Seer, have broadened the availability of low-abundance plasma proteins, enhancing proteome coverage (11). Despite these advancements, plasma remains one of the most challenging biofluids for proteomic analysis because of its extreme dynamic range and the dominance of highly abundant proteins (12, 13). Hence, additional strategies that can enhance proteome coverage while maintaining analytical simplicity and throughput are still highly desirable.

To address this challenge, we previously developed tomato lectin affinity purification-based mass spectrometry (TomAP-MS), a workflow that improves plasma proteome coverage via tomato lectin-mediated enrichment (14). By utilizing the affinity of tomato lectin for specific glycan structures, this strategy enables deeper access to plasma proteins that are challenging to observe in unfractionated samples, offering a valuable framework for biomarker-focused plasma analysis. In the original TomAP-MS workflow, bound proteins were eluted using a buffer containing 4% sodium dodecyl sulfate (SDS), followed by SP3-based cleanup and digestion. While effective, this approach involved multiple handling steps, heightened workflow complexity, and limited sample throughput, diminishing its practicality for wider analytical applications.

To simplify the workflow, we explored acid elution as an alternative to SDS-based elution, reasoning that proteins recovered under acidic conditions could be directly subjected to tryptic digestion after pH adjustment. Conventional acid elution often results in reduced protein recovery. To address this limitation, we investigated the use of lauryl maltose neopentyl glycol (LMNG), a surfactant that is compatible with tryptic digestion and can be readily removed using reversed-phase StageTips (15). In this study, we developed an improved TomAP-MS workflow incorporating LMNG-assisted acid elution (LAcE) and evaluated its performance in comparison with conventional acid elution and the original SDS/SP3 workflow. The broader applicability and practical utility of this approach in plasma proteomics and related workflows were also evaluated.

## Materials and Methods

### Preparation of Plasma and Serum

Peripheral blood was collected from healthy adult volunteers. Ethylenediaminetetraacetic acid (EDTA) plasma and heparin plasma were prepared using EDTA-treated (cat. no. VP-DK050K, Terumo Corporation, Tokyo, Japan) and heparin-treated blood collection tubes (cat. no. VO-H050K, Terumo Corporation), respectively, and the samples were centrifuged promptly after blood collection. Serum was prepared using serum collection tubes (cat. no. VP-AS109K50, Terumo Corporation). Blood collected for serum preparation was allowed to clot at room temperature for 30 min before centrifugation. After centrifugation, the plasma or serum supernatants were stored at −80 °C until analysis. Additionally, 200 EDTA plasma samples from healthy donors were purchased from Central Link Inc. (Tokyo, Japan) and stored at −80 °C until use.

This study was conducted following the Ethical Guidelines for Medical and Biological Research Involving Human Subjects and approved by the Institutional Ethical Committee of Kazusa DNA Research Institute (approval no. 2020-06 and 2025-01).

### TomAP-SDS/SP3 workflow (conventional workflow)

A 25 µL aliquot of streptavidin bead suspension (cat. no. 21152104010150, Cytiva, Marlborough, MA, USA) was added to 600 µL of protein-free blocking buffer (Setsuyaku-Kun Supporter, DRC, Tokyo, Japan) diluted 10-fold with Tris-buffered saline (TBS; 25 mM Tris-HCl, pH 7.4, 137 mM NaCl, and 2.68 mM KCl). Subsequently, 10 µL of 2 µg/µL biotinylated tomato lectin (cat. no. B-1175-1, Vector Laboratories, Burlingame, CA, USA) was added. The mixture was gently agitated for 30 min, and the beads were washed once with 1.2 mL of dilution/wash (D/W) buffer (TBS containing 0.0005% Tween 20). Subsequently, 50 µL of plasma diluted in 450 µL of D/W buffer was added to the beads and mixed for 60 min. The beads were washed twice with 1.2 mL of D/W buffer and eluted by mixing for 15 min with 200 µL of elution buffer, which comprised 100 mM Tris-HCl, pH 8.0 containing 4% SDS. The eluted proteins were collected. This workflow was automated using either a Maelstrom 9610 or a Maelstrom 8 instrument (Taiwan Advanced Nanotech, Taoyuan, Taiwan).

The 200 µL eluate was cleaned up and digested using the SP3-LASP method, a low-loss SP3-based protocol previously described (15), employing either a Maelstrom 9610 or a Maelstrom 8 instrument. Briefly, two types of SeraMag SpeedBead carboxylate-modified magnetic particles (hydrophilic particles, cat. no. 45152105050250, and hydrophobic particles, cat. no. 65152105050250; Cytiva) were mixed at a 1:1 (v/v) ratio, washed twice with distilled water, and reconstituted in distilled water at a concentration of 10 µg solids/µL. Subsequently, 20 µL of the reconstituted beads were added to each sample, followed by the addition of 1-propanol to a final concentration of 75% (v/v) and mixed for 5 min. After removing the supernatant, the bead pellet was washed twice with 80% 1-propanol. The beads were resuspended in 80 µL of 50 mM Tris-HCl (pH 8.0), 10 mM CaCl_2_, and 0.02% LMNG, and digested with 1 µg of Trypsin/Lys-C Mix (cat. no. V5072, Promega, Madison, WI, USA) at 37 °C for 16 h. The peptides obtained were reduced and alkylated by adding 8 µL of a solution containing 110 mM tris(2-carboxyethyl) phosphine and 440 mM 2-chloroacetamide, followed by incubation at 80 °C for 15 min. The samples were acidified with 16 µL of 5% trifluoroacetic acid (TFA). Peptides were desalted using a GL-Tip SDB (GL Sciences, Tokyo, Japan) following the manufacturer’s instructions. Subsequently, the peptides were eluted with 34% acetonitrile in 0.1% TFA and dried using a centrifugal evaporator (miVac Duo Concentrator, Genevac). The dried peptides were reconstituted in 0.02% DMNG containing 0.1% TFA.

### TomAP-LAcE workflow (optimized workflow)

A 25 µL aliquot of streptavidin bead suspension (cat. no. 21152104010150, Cytiva) was added to 600 µL of protein-free blocking buffer (Setsuyaku-Kun Supporter, DRC), diluted 10-fold with TBS. Subsequently, 10 µL of 2 µg/µL biotinylated tomato lectin (cat. no. B-1175-1, Vector Laboratories) was added. The mixture was gently agitated for 30 min, and the beads were washed once with 1.2 mL of D/W buffer. Subsequently, 50 µL of plasma or serum, diluted in 450 µL of D/W buffer, was added to the beads and mixed for 60 min. In low-input experiments, 1 or 5 µL of plasma was diluted in 100 µL of D/W buffer and then incubated with the beads under identical conditions. The beads were subsequently washed twice with 1.2 mL of D/W buffer. The beads were subsequently rinsed with 1.2 mL of TBS to remove residual Tween 20 carried over from the D/W buffer. For protein elution, the beads were mixed with 70 µL of elution solvent for 15 min. The solvents tested included 0.5% TFA, 0.05% LMNG, and combinations of 0.05% LMNG with 0.5% TFA. The optimization results led to the adoption of 0.5% TFA containing 0.05% LMNG as the standard LAcE condition for subsequent experiments. This workflow was performed automatically using either a Maelstrom 9610 or a Maelstrom 8 instrument (Taiwan Advanced Nanotech).

Following magnetic separation, the eluate was collected and neutralized by adding 500 mM Tris-HCl, pH 8.0, with 20 mM CaCl_2_ to adjust the pH to approximately 8.0. For digestion, 1 µg of Trypsin/Lys-C Mix (cat. no. V5072, Promega) was added to the neutralized eluate and then incubated at 37 °C for 16 h. The peptides obtained were reduced and alkylated by adding 14 µL of a solution with 110 mM tris(2-carboxyethyl) phosphine and 440 mM 2-chloroacetamide, then incubated at 80 °C for 15 min. Subsequently, the samples were acidified with 35 µL of 5% TFA. Peptides were desalted using a GL-Tip SDB (GL Sciences) following the manufacturer’s instructions, eluted with 34% acetonitrile in 0.1% TFA, and dried using a centrifugal evaporator (miVac Duo Concentrator, Genevac). The dried peptides were reconstituted in 0.02% DMNG containing 0.1% TFA.

### Purification of extracellular vesicle (EVs) from human serum

Human serum extracellular vesicles (EVs) were isolated using the Tim4-phosphatidylserine (PS) affinity method combined with the MagCapture Exosome Isolation Kit PS Ver.2 (Wako Pure Chemical) and a Maelstrom 8 instrument (Taiwan Advanced Nanotech). Briefly, 150 μL of healthy human serum was centrifuged at 3000 g at 4 °C for 20 min, and then, 100 μL of the supernatant was aliquoted into separate tubes. Subsequently, 300 μL of TBS and 1 μL of exosome binding enhancer (provided with the kit) were added to the supernatants and gently mixed. To prepare the beads for the Tim4-PS affinity method, 30 μL of exosome capture beads (provided with the kit) were washed once with 250 μL of exosome immobilizing/washing buffer (provided with the kit). Subsequently, the beads were mixed in 10 μL of biotin-labeled exosome capture (diluted by 250 μL of exosome immobilizing/washing buffer) and agitated at 1000 rpm for 10 min. After washing twice with 250 μL of exosome immobilizing/washing buffer, the beads were added to the samples. To bind EVs to the beads, the mixture was agitated at 1000 rpm for 2 h, then washed twice with 250 μL of exosome immobilizing/washing buffer. In the LAcE workflow, the beads were further rinsed with 1.2 mL of TBS to eliminate any remaining exosome immobilization/wash buffer from the prior step (this step was excluded in the SDS/SP3 workflow). Finally, bead-captured EVs were eluted by mixing for 15 min with elution solvent (200 μL of 100 mM Tris–HCl pH 8.0 containing 4% SDS for the SDS/SP3 workflow or 70 μL of 0.5% TFA containing 0.05% LMNG for the LAcE workflow).

Samples eluted with 100 mM Tris-HCl (pH 8.0) containing 4% SDS were processed using the same SP3 procedure as in the TomAP-SDS/SP3 workflow, whereas samples eluted with 0.5% TFA containing 0.05% LMNG were digested in the same manner as in the TomAP-LAcE workflow.

### Analysis of human RAGE-binding proteins

A 5 µL aliquot of streptavidin bead suspension (cat. no. 21152104010150, Cytiva) was added to 600 µL of radioimmunoprecipitation assay buffer (cat. no. 89901, Thermo Fisher Scientific, Waltham, MA, USA). Subsequently, 16 µL of 250 ng/µL biotinylated human RAGE (cat. no. 11629-H49H-B, Sino Biological, Beijing, China) was added. The mixture was gently agitated for 30 min, and the beads were washed once with 1.2 mL of D/W buffer. Subsequently, 50 µL of plasma diluted in 450 µL of D/W buffer was added to the beads and mixed for 60 min. The beads were subsequently rinsed twice with 1.2 mL of D/W buffer.

In the LAcE workflow, the beads were rinsed with 1.2 mL of TBS to remove residual Tween 20 from the D/W buffer. This step was omitted in the SDS/SP3 workflow. Finally, bead-captured proteins were eluted by mixing for 15 min with appropriate elution solvent: 200 μL of 100 mM Tris–HCl (pH 8.0) containing 4% SDS for the SDS/SP3 workflow, or 70 μL of 0.5% TFA containing 0.05% LMNG for the LAcE workflow.

Samples eluted with 100 mM Tris-HCl (pH 8.0) containing 4% SDS were processed using the same SP3 procedure as in the TomAP-SDS/SP3 workflow, whereas samples eluted with 0.5% TFA containing 0.05% LMNG were digested in the same manner as in the TomAP-LAcE workflow.

### NanoLC-MS/MS

NanoLC-MS/MS measurements were performed using three methods with different LC programs: an 84.5-min gradient (16-SPD), a 54-min gradient (24-SPD), and a 23-min gradient (50-SPD). An aliquot of the redissolved peptides equivalent to 20 µL of the plasma input was directly injected onto a 75 µm × 30 cm nanoLC column (ReproSil-Pur C18, 1.5 µm particle size, 100 Å pore size; CoAnn Technologies, Richland, Washington, USA) maintained at 60 °C. Mobile phase A consisted of 0.1% formic acid in distilled water, and mobile phase B consisted of 0.1% formic acid in 80% acetonitrile. For the 16-SPD method, peptides were separated using an 84.5-min gradient as follows: 1% B at a flow rate of 600 nL/min for 0–0.5 min; 1–6% B at 600–200 nL/min for 0.5–4.5 min; 6–24% B at 200 nL/min for 4.5–60 min; 24–40% B at 200 nL/min for 60–78.5 min; 40–98% B at 200 nL/min for 78.5–79.5 min; 98% B at 200 nL/min for 79.5–80.5 min; 98% B at 200–600 nL/min for 80.5–81.5 min; 98% B at 600–750 nL/min for 81.5–82.5 min; and 98% B at 750 nL/min for 82.5–84.5 min. For the 24-SPD method, peptides were separated using a 54-min gradient as follows: 1% B at 700 nL/min for 0–0.5 min; 1–7% B at 700–250 nL/min for 0.5–3.5 min; 7–26% B at 250 nL/min for 3.5–38.5 min; 26–42% B at 250 nL/min for 38.5–49.5 min; 42–98% B at 250 nL/min for 49.5–50.5 min; 98% B at 250 nL/min for 50.5–52 min; 98% B at 250–800 nL/min for 52–53.5 min; and 98% B at 800 nL/min for 53.5–54 min. For the 50-SPD method, peptides were separated using a 23-min gradient as follows: 1% B at 700 nL/min for 0–0.5 min; 1–10% B at 700–400 nL/min for 0.5–3.5 min; 10–28% B at 400 nL/min for 3.5–15.5 min; 28–42% B at 400 nL/min for 15.5–19.5 min; 48–98% B at 400 nL/min for 19.5–20.5 min; 98% B at 400 nL/min for 20.5–21.5 min; 98% B at 400–800 nL/min for 21.5–22.5 min; and 98% B at 800 nL/min for 22.5–23 min.

Eluted peptides were analyzed using an Orbitrap Astral mass spectrometer (Thermo Fisher Scientific) equipped with an InSpIon system (16). MS1 spectra were acquired over an m/z range of 380–980 at a resolution of 240,000 using the Orbitrap analyzer, with an AGC target of 300% and a maximum injection time of 5 ms. MS2 spectra were acquired over an m/z range of 200–2,000 using the Orbitrap Astral analyzer. The AGC target was set to 300% for 16-SPD and 500% for 24 and 50-SPD, with maximum injection times of 3.5 ms for 16-SPD and 3 ms for 24- and 50-SPD. A normalized collision energy of 25% was applied, and the isolation width was set to 2 Th for 16 and 24-SPD and 3 Th for 50-SPD, with optimized window placement.

### Data analysis

The data-independent acquisition mass spectrometry (DIA-MS) data were searched against an in silico human spectral library using DIA-NN v2.5.0 Enterprise (https://github.com/vdemichev/DiaNN) (17). The spectral library was initially generated in DIA-NN from the UniProt human proteome database (proteome ID: UP000005640; 20,659 entries; downloaded on April 1, 2026). The spectral library generation parameters included the following: protease, Trypsin/P; misses, 1; peptide length, 7–45; precursor charge, 2–4; precursor m/z, 380–980; and fragment m/z, 200-2,000. Additionally, “Predicting from FASTA,” “N-term M excision,” and “Carbamidomethyl (C)” were enabled. For the DIA-NN search, the following parameters were applied: MS1 accuracy, 4 ppm; MS2 accuracy, 8 ppm; scoring, generic; proteotypicity, isoform IDs; machine learning, NNs (cross-validated); and Quantification strategy, QuantUMS (precision). Furthermore, “Unrelated runs,” “Protein inference,” and “Knowledge base” were enabled. For protein identification analysis, each dataset was individually analyzed, and the number of identified proteins was tallied for each run. For quantitative comparisons between methods, the datasets were analyzed with MBR enabled and cross-run normalization disabled. The 200 plasma samples from healthy individuals were analyzed using MBR with cross-run normalization set to RT-dependent. The false discovery rate threshold was set to 1% or less at both the precursor and protein levels. Protein quantification values were obtained by aggregating the quantitative values of unique peptides calculated using DIA-NN. Statistical significance for two-group comparisons was evaluated using a two-sided Welch’s t-test. Statistical significance is denoted as follows: *P < 0.05, **P < 0.005, and ***P < 0.0005. Pearson’s correlation analysis was conducted using R, and the figures were created using ggplot2. Additional graphs were created using GraphPad Prism 9 (GraphPad Software, San Diego, CA, USA). Proteins associated with OMIM were identified using the OMIM_DISEASE annotation category in DAVID (https://david.ncifcrf.gov/tools.jsp). To evaluate relevance to newborn screening, the gene list corresponding to the identified proteins was compared with the list of genes associated with conditions included in the Recommended Uniform Screening Panel (RUSP), and proteins mapped to matched genes were defined as RUSP-associated proteins.

## Results and Discussion

### Development of the LAcE workflow for TomAP-MS

To streamline the TomAP-MS workflow, we evaluated acid elution as an alternative to the original SDS-based elution method (Figure 1). Because proteins recovered by acid elution can be directly subjected to tryptic digestion after pH adjustment, this strategy eliminates the need for SP3 cleanup and offers a simpler workflow. However, applying conventional acid elution to TomAP-MS reduced the number of identified proteins compared to the original SDS/SP3 workflow (Figure 2A). These results suggest that while acid elution enhances procedural simplicity, its typical use in TomAP-MS compromises proteome coverage.

**Figure 1.**
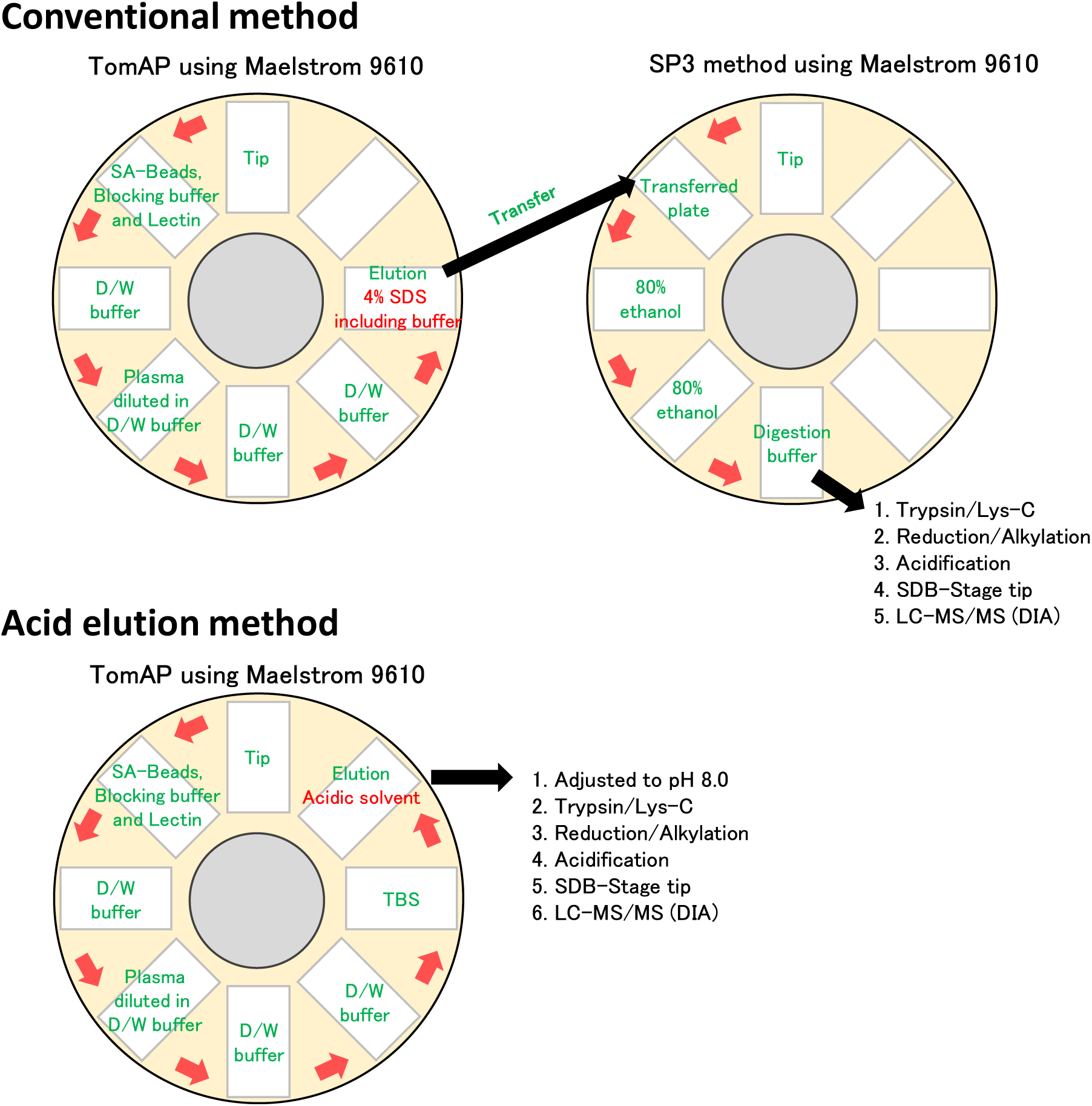
Automated method for enrichment of trace plasma proteins by tomato lectin affinity purification (TomAP) using the Maelstrom 9610 instrument. The Maelstrom 9610 instrument is equipped to accommodate eight different 96-deep-well plates, one of which requires specific tip placement for proper operation. Each plate can be positioned beneath a 96-pin magnetic head. This head descends into the 96-well plate, enabling binding, release, or mixing of magnetic beads in solution. For the conventional TomAP method, the Maelstrom 9610 instrument was used to automatically process the reaction of biotinylated lectin with streptavidin (SA) beads, the interaction of tomato lectin-bound beads with plasma, the washing steps between each stage, and the elution step. Proteins bound to the beads were eluted by mixing for 15 min in a buffer containing sodium dodecyl sulfate (SDS). The lectin-enriched plasma was automatically processed using the Maelstrom 9610 instrument with single-pot, solid-phase-enhanced sample preparation (SP3), followed by Trypsin/Lys-C digestion, reduction and alkylation, and desalting using StageTips. For the acid-elution TomAP method, the beads were rinsed with TBS prior to elution to prevent solvent carryover from previous steps, and proteins were eluted with an acidic solvent instead of the 4% SDS-containing buffer. After elution, the samples were adjusted to approximately pH 8.0, followed by Trypsin/Lys-C digestion, reduction and alkylation, and desalting using StageTips.

**Figure 2.**
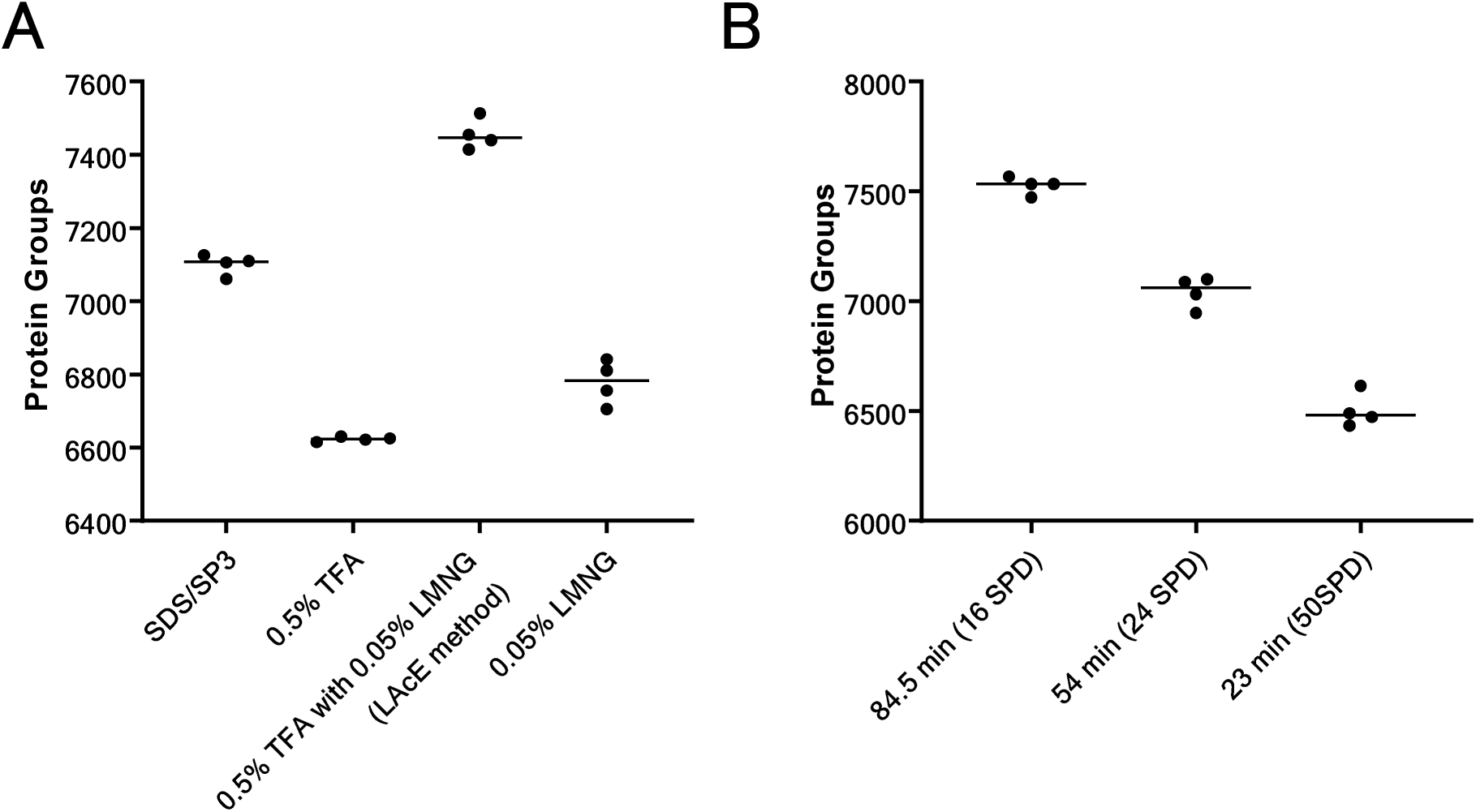
Comparison of elution conditions and LC–MS throughput settings for the TomAP-LAcE workflow. (A) Comparison of the number of protein groups identified using the SDS/SP3 workflow, acid elution with 0.5% trifluoroacetic acid (TFA), lauryl maltose neopentyl glycol (LMNG)-assisted acid elution with 0.5% TFA and 0.05% LMNG (LAcE), and 0.05% LMNG alone. (B) Number of protein groups identified from EDTA plasma processed using the TomAP-LAcE workflow and analyzed using liquid chromatography–mass spectrometry (LC–MS) with different gradient lengths: 84.5 min (16-SPD), 54 min (24-SPD), and 23 min (50-SPD). Each dot represents an independent analysis (n = 4 per group). Protein group identifications were obtained by analyzing each raw data file individually. Horizontal bars indicate median values.

To address this reduction, we explored the use of a surfactant that is compatible with tryptic digestion and readily removable by reversed-phase StageTips. We focused on LMNG, which does not substantially interfere with tryptic digestion and can be efficiently removed by reversed-phase StageTips (15). Based on this rationale, we established an improved TomAP-MS workflow incorporating LMNG-assisted acid elution (LAcE). In this workflow, proteins were eluted under acidic conditions in the presence of LMNG, followed by pH adjustment and direct tryptic digestion without SP3 cleanup. This modified procedure reduced the complexity of sample handling and improved throughput.

We then compared the analytical performance of the LAcE workflow with that of conventional acid elution and the original SDS/SP3 workflow. The LAcE workflow yielded a greater number of protein identifications compared with that of conventional acid elution (0.5% TFA), suggesting that the addition of LMNG effectively offset the protein loss linked to acid elution alone. Notably, LAcE also outperformed the original SDS/SP3 workflow, demonstrating that simplification of the elution and digestion procedure did not compromise proteome depth. The combination of LMNG and acid elution enhanced analytical performance and streamlined the workflow.

Analyzing human plasma processed using this optimized workflow via LC-MS revealed the identification of up to 7,567 proteins with an 84.5-min gradient (about 16 samples/day), 7,099 proteins with a 55-min gradient (about 24 samples/day), and 6,615 proteins with a 23-min gradient (about 50 samples/day) (Figure 2B and Table S1). To our knowledge, one of the highest numbers reported in deep plasma proteome analyses by LC-MS under comparable analysis times is 5,163 proteins from plasma processed by the Mag-Net method with a 60-min gradient (18). The detection of over 7,000 proteins within a 54-min analysis, which is broadly similar in duration, underscores the efficient performance of the optimized TomAP workflow. To evaluate the wider applicability of this approach, we investigated if LAcE could be utilized in proteomic workflows beyond TomAP-MS (Figure 3A, B). We applied LAcE to Tim4-based EV enrichment and protein interaction analysis workflows. In EV enrichment, LAcE increased the number of identified proteins compared with that of the conventional SDS/SP3 workflow and enhanced the recovery of known EV markers CD9, CD81, TSG101, PDCD6IP, SDCBP, FLOT1, and FLOT2 (19, 20). However, the retrieval of CD63 decreased. The decrease may be owing to factors like instability under acidic conditions; however, the precise cause remains undetermined. Altogether, these results suggest that the LAcE workflow offers a general advantage for EV enrichment.

**Figure 3.**
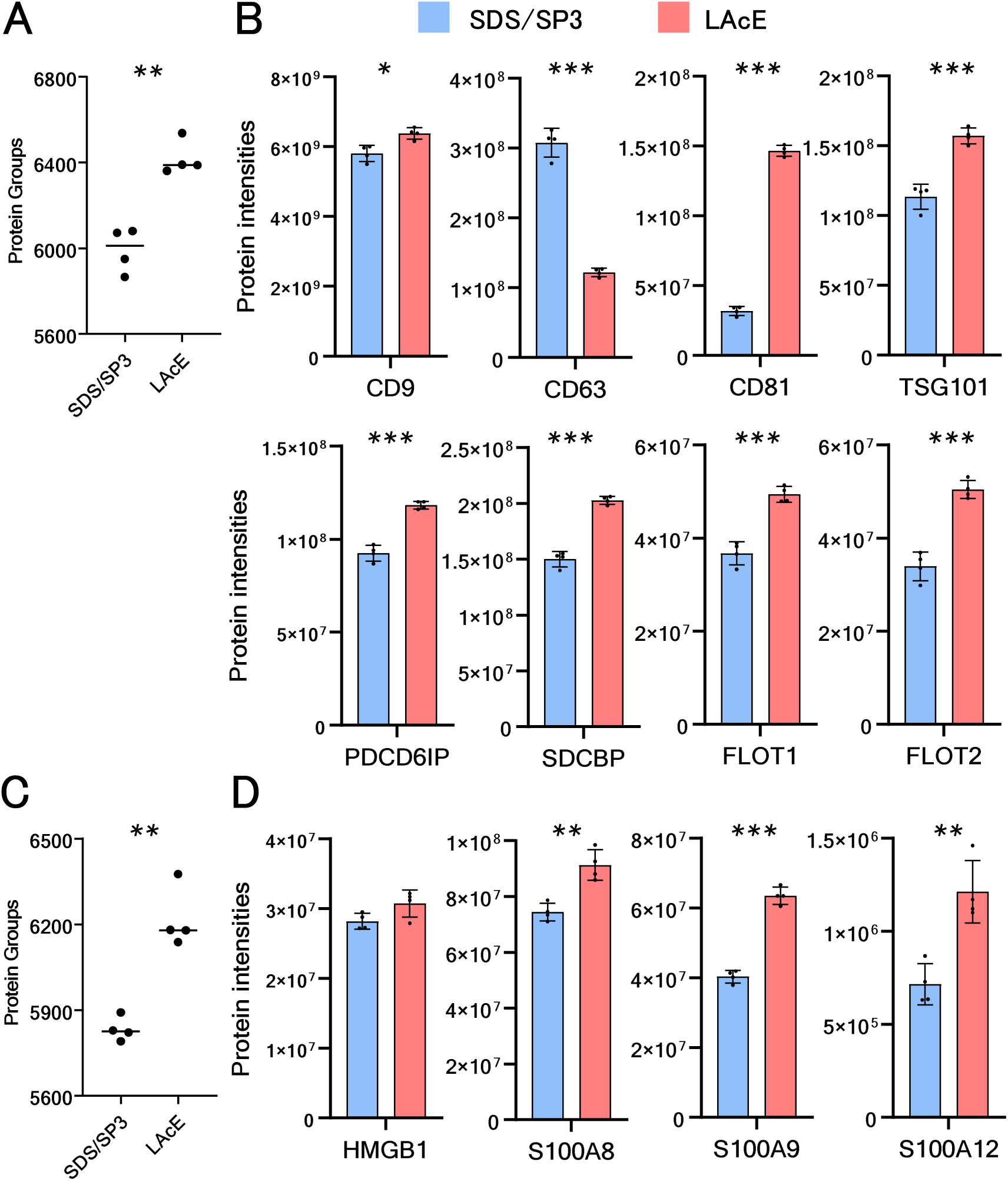
Application of the LAcE workflow to extracellular vesicle enrichment and protein interaction analysis workflows. (A) Comparison of the number of protein groups identified from human serum extracellular vesicles (EVs) enriched by the Tim4–phosphatidylserine affinity method and processed using either the conventional SDS/SP3 workflow or the lauryl maltose neopentyl glycol-assisted acid elution (LAcE) workflow. (B) Protein intensities of representative EV marker proteins, including CD9, CD63, CD81, TSG101, PDCD6IP, SDCBP, FLOT1, and FLOT2, comparing the SDS/SP3 and LAcE workflows. (C) Comparison of the number of protein groups identified in AP-MS analysis of human RAGE-binding proteins using the SDS/SP3 and LAcE workflows. (D) Protein intensities of representative known human RAGE interactors, including HMGB1, S100A8, S100A9, and S100A12, comparing the SDS/SP3 and LAcE workflows. Each dot represents an independent analysis (n = 4 per group). Horizontal bars in panels A and C indicate median values. Protein group identifications were obtained by analyzing each raw data file individually, whereas protein intensities were obtained from joint analysis of the datasets being compared. Bars indicate mean values, and error bars indicate standard deviation (SD). Statistical significance was assessed using a two-sided Welch’s t-test and is indicated as follows: *P < 0.05, **P < 0.005, and ***P < 0.0005.

For protein-protein interaction analysis, we conducted affinity purification-mass spectrometry (AP-MS) targeting human RAGE (Figure 3C, D). In this analysis, LAcE increased the number of identified proteins compared with that of the conventional SDS/SP3 workflow. It also improved the recovery of the known human RAGE interactors HMGB1, S100A8, S100A9, and S100A12 (21–24). However, an increasing trend was observed only for HMGB1, with no statistically significant difference. These observations imply that the benefits of LAcE are not limited to TomAP-MS and may apply to other proteomic workflows needing effective protein elution.

Overall, the LAcE workflow reduces the need for detergent-removal and SP3-based cleanup prior to digestion, thereby decreasing sample-handling steps and improving suitability for high-throughput processing. Collectively, these results indicate that LAcE provides a practical improvement to TomAP-MS by increasing protein identifications while preserving the operational advantages of a simpler and faster workflow.

### Specimen type, low-input performance, and freeze–thaw tolerance in the optimized TomAP-MS workflow

We then evaluated practical factors influencing the performance of TomAP-MS in plasma proteomics, such as specimen type, compatibility with low input, and tolerance to repeated freeze–thaw cycles. Blood specimen type and pre-analytical handling significantly affect circulating proteome measurements, making them crucial considerations in plasma proteomics workflows (3, 25). Among EDTA plasma, heparin plasma, and serum, EDTA plasma produced the most protein identifications, followed by heparin plasma and serum (Figure 4A). These results indicate that specimen type significantly affects proteome coverage in TomAP-MS, and suggest that EDTA plasma is the most suitable specimen for biomarker discovery applications requiring broad protein identification.

**Figure 4.**
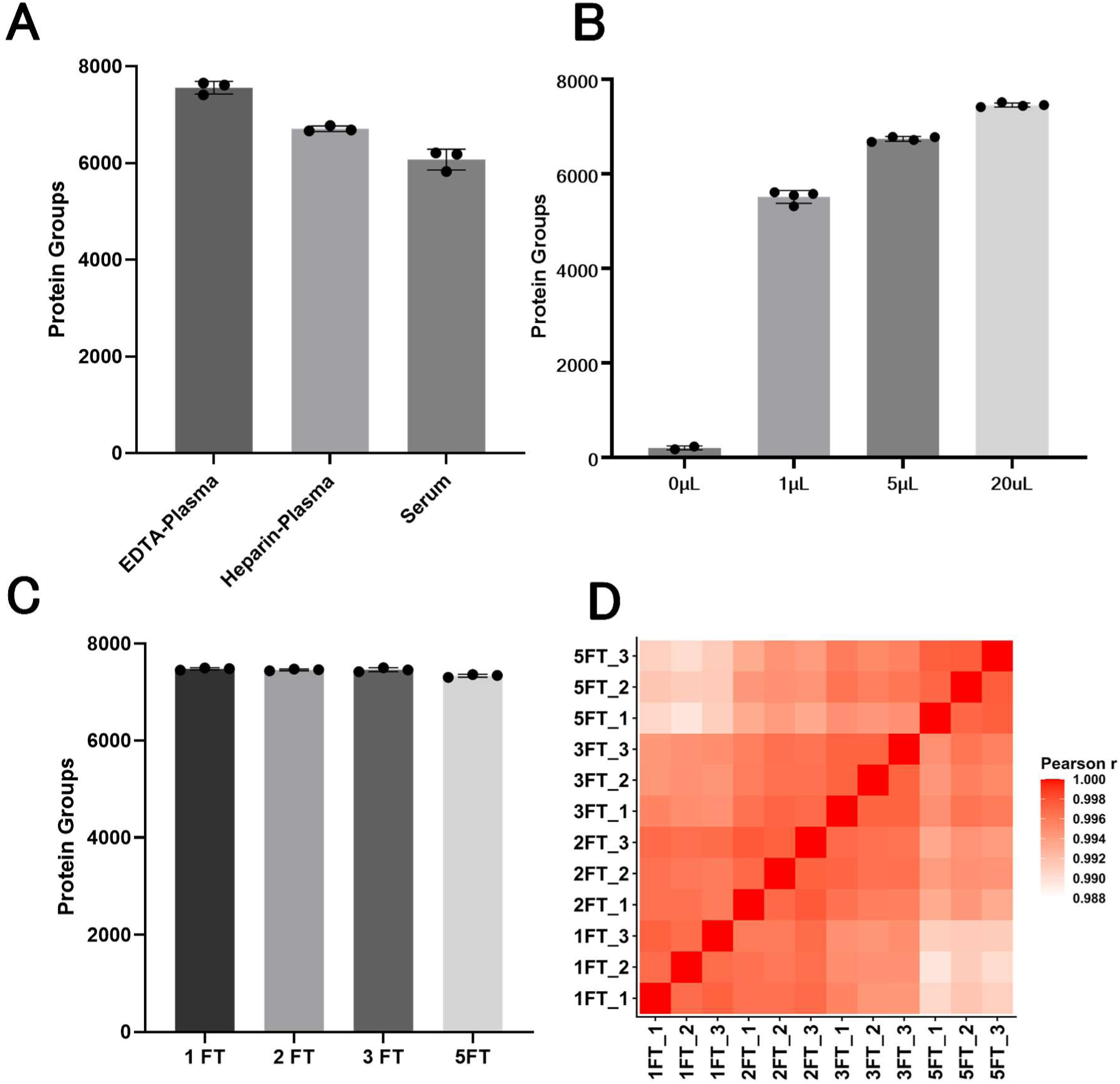
Evaluation of specimen type, low-input performance, and freeze–thaw tolerance in the TomAP-LAcE workflow. (A) Number of protein groups identified from different blood specimen types, including ethylenediaminetetraacetic acid (EDTA) plasma, heparin plasma, and serum, analyzed using the tomato lectin affinity purification–lauryl maltose neopentyl glycol-assisted acid elution (TomAP-LAcE) workflow. (B) Number of protein groups identified from different input volumes of EDTA plasma (0, 1, 5, and 20 µL), demonstrating the low-input compatibility of the workflow. (C) Number of protein groups identified from EDTA plasma samples subjected to repeated freeze–thaw (FT) cycles (1, 2, 3, and 5 cycles). (D) Pearson correlation matrix of proteomic profiles obtained from EDTA plasma samples subjected to repeated freeze–thaw cycles, showing overall similarity across replicates and freeze–thaw conditions. Each dot represents an independent analysis. Protein group identifications were obtained by analyzing each raw data file individually. The numbers of independent analyses were n = 3 for panels A, C, and D, and n = 4 for panel B, except for the “0 µL” sample, for which n = 2. Bars indicate mean values, and error bars indicate standard deviation (SD).

We then examined whether TomAP-MS could be applied to low-input plasma samples (Figure 4B). The optimized workflow using EDTA plasma enabled the identification of over 5,000 proteins from just 1 µL of input material. This result shows that TomAP-MS maintains substantial analytical depth despite minimal sample quantities, making it valuable in environments with limited sample access.

Finally, we assessed the impact of repeated freeze-thaw cycles on TomAP-MS profiles of EDTA plasma (Figure 4C, D). Repeated freeze–thaw cycles are a well-recognized pre-analytical factor that can impact blood proteome measurements (25, 26). Even after five freeze-thaw cycles, the decrease in the number of identified proteins was only approximately 2%, suggesting no substantial change. In contrast, protein intensity correlations exhibited minimal changes after one to three freeze-thaw cycles, but significant alterations were observed after five cycles. These results indicate that EDTA plasma samples subjected to up to three freeze-thaw cycles are appropriate for TomAP-MS-based proteomic analysis; however, excessive freeze-thawing should be circumvented in practical applications.

These results offer practical guidance for utilizing TomAP-MS in plasma proteomics. They endorse the use of EDTA plasma as the preferred specimen type, demonstrate the workflow’s applicability to low-input samples, and define an acceptable range of freeze-thaw handling for robust analysis.

### Large-scale plasma profiling in 200 healthy individuals for inherited disease applications

To further assess the robustness and clinical applicability of the optimized TomAP-MS workflow, we analyzed plasma samples from 200 healthy individuals employing the 50-SPD setting. The median number of identified proteins per sample was about 6,000, showing that this workflow allows consistent and extensive plasma proteome profiling even in a large cohort (Figure 5A). A total of 4,117 proteins were commonly detected across all 200 individuals, and these proteins were considered to constitute a core plasma proteome that can be reproducibly observed in healthy individuals using TomAP-MS.

**Figure 5.**
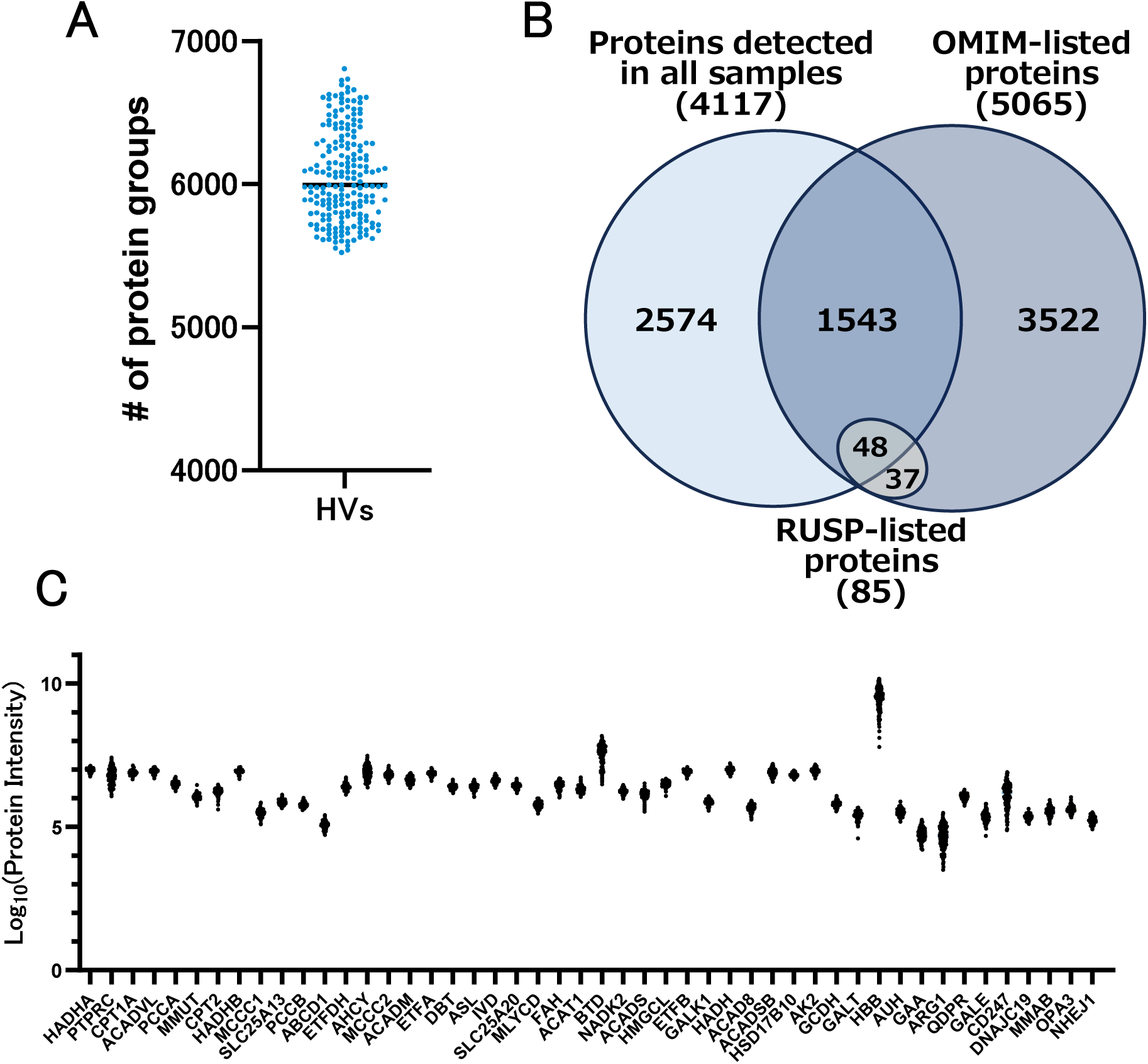
Reproducible detection of inherited disease–related proteins in plasma from 200 healthy individuals using TomAP-MS. (A) Number of protein groups identified per sample in plasma from 200 healthy volunteers (HVs) analyzed using tomato lectin affinity purification-based mass spectrometry (TomAP-MS) with the 50-SPD setting. Each dot represents one sample, and the horizontal bar indicates the median number of identified protein groups across all 200 samples (5,995). (B) Overlap of proteins detected in all samples with OMIM-listed proteins and Recommended Uniform Screening Panel (RUSP)-listed proteins. Of the 4,117 proteins commonly detected across all samples, 1,543 were annotated in OMIM and 48 corresponded to RUSP-listed proteins. (C) Protein intensity profiles of the 48 RUSP-listed proteins detected in all 200 samples. Each dot represents the log_10_ protein intensity of one sample.

We then evaluated the disease relevance of 4,117 proteins and identified that 1,543 were annotated in OMIM, a comprehensive knowledge base of human genes and genetic disorders (27, 28) (Figure 5B and Table S2). This suggests that a substantial proportion of the frequently identified proteins are gene products linked to recognized Mendelian disorders. Additionally, 48 proteins corresponded to disorders in the RUSP, the list of conditions recommended by the US Secretary of Health and Human Services for newborn screening programs (29, 30). Quantitative assessment of these 48 proteins across individual samples indicated their relatively stable abundance (Figure 5C and Table S2). These observations indicate that the proteins consistently identified using TomAP-MS encompass numerous targets directly pertinent to the pathophysiology and diagnosis of inherited disorders.

Directly detecting disease-relevant proteins may provide several advantages for inherited disease diagnostics. Protein-level analysis can capture the functional consequences of pathogenic variants, thus offering complementary information to genetic testing (31, 32). Proteomic analysis can identify decreased or missing protein expression when pathogenic changes impact gene regulation, protein stability, or degradation in ways not easily deduced from sequence data alone (31, 32). Some pathogenic variants cause disease by disrupting protein folding or stability, resulting in accelerated degradation and decreased protein levels, assessable directly at the proteome level (31). Broad proteome profiling can capture both the primary affected protein and downstream pathway perturbations, offering a functional readout for variant interpretation and disease stratification (32, 33).

There is a clear need for this approach as similar efforts have been undertaken in dried blood spot (DBS) proteomics to comprehensively evaluate inherited and rare diseases (34–37). From this perspective, the current findings indicate that TomAP-MS could broaden its applications by facilitating extensive proteomic analysis not just from DBS-derived samples but also from plasma. TomAP-MS may serve as a versatile platform for comprehensive inherited disease diagnostics across various specimen types.

Altogether, these findings support the potential utility of TomAP-MS in inherited disease diagnostics. In particular, the reproducible detection of many disease-related proteins, including OMIM-associated and RUSP-related proteins, across plasma samples from 200 healthy individuals suggests that this workflow may provide a promising foundation for future proteomics-based screening and diagnostic applications in inherited disorders.

## Conclusion

We developed an improved TomAP-MS workflow by incorporating LAcE, which simplified sample preparation while increasing protein identifications compared with both conventional acid elution and the original SDS/SP3 workflow. The optimized workflow enables deep and high-throughput plasma proteome profiling, with broad applicability to EV enrichment and protein–protein interaction analysis. Practical evaluation further showed that EDTA plasma is the preferred specimen type for TomAP-MS, that the workflow is compatible with low-input samples, and that plasma samples subjected to up to three freeze-thaw cycles remain suitable for analysis. Moreover, large-scale profiling of plasma from 200 healthy individuals showed the reproducible detection of numerous disease-relevant proteins, endorsing the potential value of TomAP-MS for upcoming proteomics-driven screening and diagnostic purposes in inherited disorders. These findings establish TomAP-MS with LAcE as a versatile and practical platform for deep plasma proteomics and related applications.

## Funding

This study was supported in part by the JSPS KAKENHI under Grant Numbers 23H02465 and 24K11013. This work was also supported by the Kazusa DNA Research Institute Foundation.

## Author contributions

Y.K. and O.O. designed the study; Y.O., H.M., and D.N. performed the research; N.U. collected blood samples from healthy volunteers; Y.O., H.M., and R.K. analyzed the data; Y.O., H.M., and Y.K. wrote the manuscript. All authors read and approved the final manuscript.

## Conflict of Interest

The authors declare that they have no conflicts of interest.

## Notes

### Competing Interest Statement

The authors have declared no competing interest.

